# Incubation of oxycodone craving is associated with CP-AMPAR upregulation in D1 and D2 receptor-expressing medium spiny neurons in nucleus accumbens core and shell

**DOI:** 10.1101/2025.04.06.647399

**Authors:** Kimberley A. Mount, Hayley M. Kuhn, Eun-Kyung Hwang, Madelyn M. Beutler, Marina E. Wolf

## Abstract

A major problem in treating opioid use disorder is persistence of craving after protracted abstinence. This has been modeled in rodents using the incubation of craving model, in which cue-induced drug seeking increases over the first weeks of abstinence from drug self-administration and then remains high for an extended period. Incubation has been reported for several opioids, including oxycodone, but little is known about underlying synaptic plasticity. In contrast, it is well established that incubation of cocaine and methamphetamine craving depends on strengthening of glutamate synapses in the nucleus accumbens (NAc) through incorporation of calcium-permeable AMPARs (CP-AMPARs). CP-AMPARs have higher conductance than the calcium-impermeable AMPARs that mediate NAc excitatory transmission in drug-naïve animals, as well as other distinct properties. Here we examined AMPAR transmission in medium spiny neurons (MSN) of NAc core and shell subregions in rats during forced abstinence from extended-access oxycodone self-administration. In early abstinence (prior to incubation), CP-AMPAR levels were low. After 17-33 days of abstinence (when incubation is stably plateaued), CP-AMPAR levels were significantly elevated in both subregions. These results explain the prior demonstration that infusion of a selective CP-AMPAR antagonist into NAc core or shell subregions prevents expression of oxycodone incubation. Then, using transgenic rats, we found CP-AMPAR upregulation on both D1 and D2 receptor-expressing MSN, which contrasts with selective upregulation on D1 MSN after cocaine and methamphetamine incubation. Overall, our results demonstrate a common role for CP-AMPAR upregulation in psychostimulant and oxycodone incubation, albeit with differences in MSN subtype-specificity.

## Introduction

In people with opioid use disorder (OUD), craving is strongly correlated with likelihood of relapse (Lueptow et al., 2020), and this vulnerability to relapse persists during protracted abstinence (Sinha, 2011). To study mechanisms that maintain opioid craving during protracted abstinence, we use a translationally relevant model, ‘incubation of craving’. In this model, cue-induced drug seeking progressively increases over the first weeks of abstinence and then remains high for additional weeks to months. In rats, incubation has been demonstrated during forced abstinence from major classes of drugs of abuse (stimulants, nicotine, alcohol, and opioids) (Pickens et al., 2011; Chow, 2025), as well as during voluntary abstinence imposed by a mutually exclusive choice between drug and non-drug rewards or by pairing drug with punishment (Fredriksson et al., 2021a; Chow, 2025). In *humans*, incubation has been demonstrated during abstinence from cocaine, methamphetamine, nicotine, and alcohol (Bedi et al., 2011; Wang et al., 2013; Li et al., 2015a; Li et al., 2016; Parvaz et al., 2016; Zhao et al., 2021). Parallel assessments have not been conducted for opioids (Chow, 2025). Nevertheless, persistence of craving during abstinence, captured by the plateau phase of the incubation model, applies across all drugs. The model’s focus on cue-induced craving is also translationally relevant. Abstinent opioid users exhibit robust motivational bias towards opioid-associated stimuli (MacLean et al., 2018) and such bias predicts heroin relapse (Marissen et al., 2006) and is correlated with craving (Field et al., 2009).

Oxycodone is a prescription opioid with high propensity for misuse that contributed significantly to the start of the opioid epidemic (Kibaly et al., 2021). Incubation of oxycodone craving in rats occurs after forced abstinence (Blackwood et al., 2019b; Blackwood et al., 2019a; Bossert et al., 2019; Salisbury et al., 2020; Altshuler et al., 2021a; Altshuler et al., 2021b; Wong et al., 2022; Olaniran et al., 2023; Alonso-Caraballo et al., 2024; Patel and Loweth, 2024) and voluntary abstinence (Fredriksson et al., 2020; Fredriksson et al., 2021b; Fredriksson et al., 2023). Studies have begun to explore circuits and mechanisms underlying oxycodone incubation (Blackwood et al., 2019b; Blackwood et al., 2019a; Bossert et al., 2019; Fredriksson et al., 2020; Salisbury et al., 2020; Altshuler et al., 2021b; Fredriksson et al., 2021b; Fredriksson et al., 2023). However, relatively few studies have focused on the role of the nucleus accumbens (NAc) (see Discussion), even though it is a critical region for motivated behavior and contributes to opioid seeking in extinction-reinstatement models and to opioid conditioned place preference (CPP) (Hearing et al., 2018; Hearing, 2019; Reiner et al., 2019; Heinsbroek et al., 2021). Furthermore, clinical studies of abstinent opioid users indicate a significant role for the NAc (Ieong and Yuan, 2017; Welsch et al., 2020). For example, abstinent heroin users showed alterations in NAc resting state connectivity, and for some measures this depended on the duration of abstinence (Zou et al., 2015). Furthermore, in an fMRI study of heroin-dependent persons under methadone maintenance, heroin cues elicited greater craving in those who relapsed within the next 3 months compared to non-relapsers; this correlated with greater cue-induced NAc activation (Li et al., 2015b).

AMPARs are the main source of excitatory drive to medium spiny neurons (MSN), the principal neurons of the NAc (Pennartz et al., 1990; Hu and White, 1996). In NAc MSN of drug-naïve rats, nearly all AMPARs contain the GluA2 subunit and are therefore Ca^2+^-impermeable or CI-AMPARs (Kourrich et al., 2007; Conrad et al., 2008). AMPARs lacking the GluA2 subunit (CP-AMPARs) have higher conductance than CI-AMPARs, so their incorporation strengthens synapses (Isaac et al., 2007). CP-AMPARs also affect the induction of subsequent plasticity (e.g., (Mameli et al., 2011)). Thus CP-AMPAR plasticity is of high functional significance. For the psychostimulants cocaine and methamphetamine, incubation of craving ultimately depends on CP-AMPAR upregulation in NAc MSN synapses (Wolf, 2025) (see Discussion).

We previously reported that infusing a selective CP-AMPAR antagonist into either NAc core or shell prevented the expression of oxycodone incubation after 15-30 days of forced abstinence from oxycodone self-administration but did not affect oxycodone seeking in early abstinence, suggesting CP-AMPARs accumulated as abstinence progressed (Wong et al., 2022). The goal of this study was to use electrophysiological methods to definitively test for CP-AMPAR plasticity in the NAc during incubation of oxycodone craving. First, in recordings from unidentified medium spiny neurons (MSN), we demonstrated CP-AMPAR upregulation in excitatory synapses of core and shell MSN after oxycodone incubation, paralleling psychostimulant findings. Then, using transgenic rats, we demonstrated CP-AMPAR upregulation on both dopamine D1 receptor-expressing MSN (D1 MSN) and D2 receptor-expressing MSN (D2 MSN) after oxycodone incubation. This is a significant difference from psychostimulant incubation in rats, where CP-AMPAR upregulation is confined to D1 MSN (see Discussion). Our study is the first to investigate plasticity of excitatory synaptic transmission after incubation of craving for Oxy or any other opioid.

## Methods

### Subjects

Subjects were adult male (250-275 g) and female (225-250 g) wild-type rats or transgenic (TG) Long Evans rats (∼8-10 weeks old at the start of the experiment). Breeders for transgenic lines were obtained from the Rat Resource & Research Center (RRRC) and wildtype breeders from Charles River (Crl:LE, strain 006). Some experimental WT rats were obtained directly from Charles River. To distinguish D1 and D2 receptor-expressing MSN, we used knock-in rat lines, generated by CRISPR-Cas9 gene editing, that encode iCre recombinase immediately after the *Drd1a* (D1R) or *Adora2a* (adenosine 2a receptor; A2a) loci [(Pettibone et al., 2019); RRRC #856 and #857, respectively]. A2a was targeted because it is selectively expressed in D2 MSN, while the D2R is also expressed on cholinergic interneurons and other elements (Alcantara et al., 2003). Validation studies (whole genome sequencing, FISH, and Cre-driven viral expression) confirmed correctly targeted Cre expression, while behavioral studies confirmed normal learning and motivation (Pettibone et al., 2019). We also confirmed Cre targeting with RNAscope (Kawa et al., 2024). To visualize D1-positive and A2a-positive MSN for slice recordings, D1-Cre or A2a-Cre rats were bred to reporter lines expressing either ZsGreen (RRRC #797) or TdTomato (RRRC #938) in the presence of Cre. Rats were housed 3-4 per cage under a reverse 12-hour light-dark cycle with food and water provided ad libitum. All procedures were approved by the Institutional Animal Care and Use Committee of Oregon Health & Science University.

### Cather surgery

For intravenous catheter surgery, rats were anesthetized with isoflurane (MWI Animal Health, Boise, ID). A silastic tubing (The Dow Chemical Company, Midland, MI) catheter (Plastics One, Roanoke, VA) was inserted into the right or left jugular vein. Meloxicam (5 mg/kg, s.c.; Covetrus, Portland, ME) was administered as an analgesic pre- and post-operatively, and the rats were given 5-7 days to recover. During recovery, catheters were flushed every 24 hours with 0.9% sterile saline (Baxter International, Deerfield, IL) containing cefazolin (10 mg, i.v.; Covetrus, Portland, ME). During self-administration training, catheters were flushed immediately after removing rats from operant boxes. Catheter patency was confirmed by injection of sodium brevital (3-4 mg/kg) through the catheter. Rats were single housed with enrichment (nylabones and nesting packs) after surgery.

### Drug self-administration

After ∼1 week of recovery from catheter surgery, self-administration training began. Sessions started within 30 minutes after the beginning of the dark cycle. Rats self-administered oxycodone (dissolved in 0.9% saline) for 6 h/day in 10 consecutive daily sessions at a dose of 0.1 mg/kg/infusion (0.065 mL/infusion) under a fixed-ratio-1 reinforcement schedule. This dose has been used in many recent oxycodone incubation studies (Blackwood et al., 2019b; Blackwood et al., 2019a; Fredriksson et al., 2020; Altshuler et al., 2021a; Altshuler et al., 2021b; Fredriksson et al., 2021b; Fredriksson et al., 2023; Olaniran et al., 2023; Patel and Loweth, 2024). Control rats self-administered saline (0.065 mL/infusion) under an identical schedule. Self-administration training occurred in operant chambers equipped with two nose-poke holes. Nose pokes in the inactive hole had no consequence. Active hole responses activated the infusion pump and led to the presentation of a light cue (yellow light illuminated the active hole) and a timeout period (concurrent with cue presentation) during which nose pokes were recorded but did not induce drug delivery. For rats depicted in Figs. 1-3, some had a 20-sec light cue while others had a 4-sec light cue; incubation of similar magnitude was observed under both conditions (Fig. S1). Rats depicted in other figures had a 4-sec light cue. Rats remained single housed in their home cages during forced abstinence.

**Figure 1.**
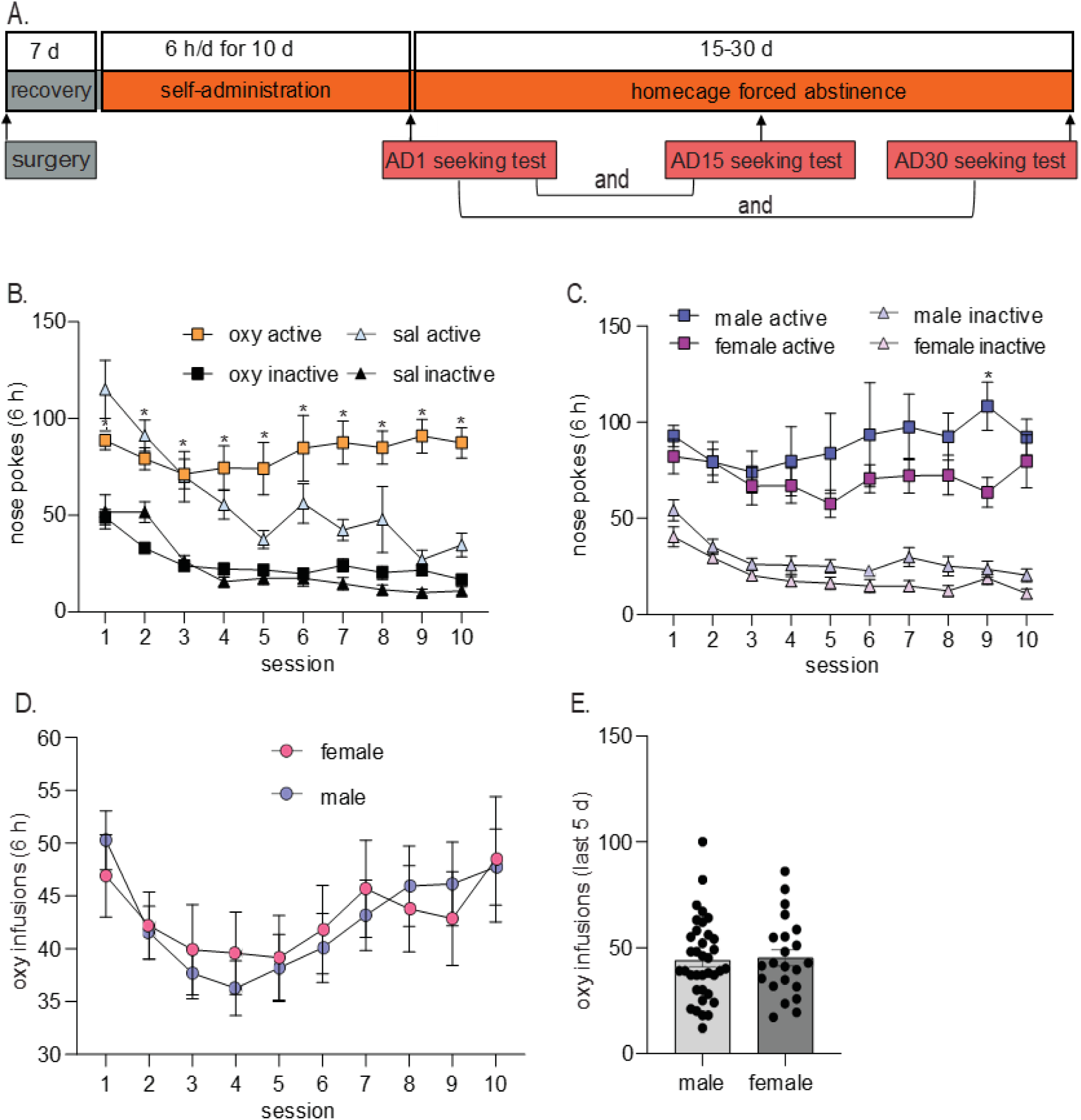
Oxycodone self-administration in male and female Long-Evans rats. (A) Rats were trained to self-administer oxycodone (oxy) or saline (sal) for 6 h/d for 10 d. Each rat underwent two cue-induced seeking tests, either on abstinence day (AD)1 and AD15 or on AD1 and AD30. (B) Average nose pokes during self-administration training for oxycodone and saline rats (\**p<*0.05 compared to inactive hole nose pokes during oxycodone self-administration). (C) Active and inactive nose poke activity across oxycodone self-administration in male and female rats (\**p*<0.05 active male vs. active female). (D) Number of infusions across all self-administration sessions for male and female rats. No sex differences in oxycodone infusions were detected on any training day. (E) Total number of oxycodone infusions averaged over the last 5 days of self-administration did not differ between males and females. Data are presented as mean ± SEM, with individual data points shown on bar graphs. Saline rats, *n=*15 (9 males, 6 females). Oxycodone rats, *n=*58 (36 males, 22 females). More information about statistical analysis is presented in the Results section.

**Figure 2.**
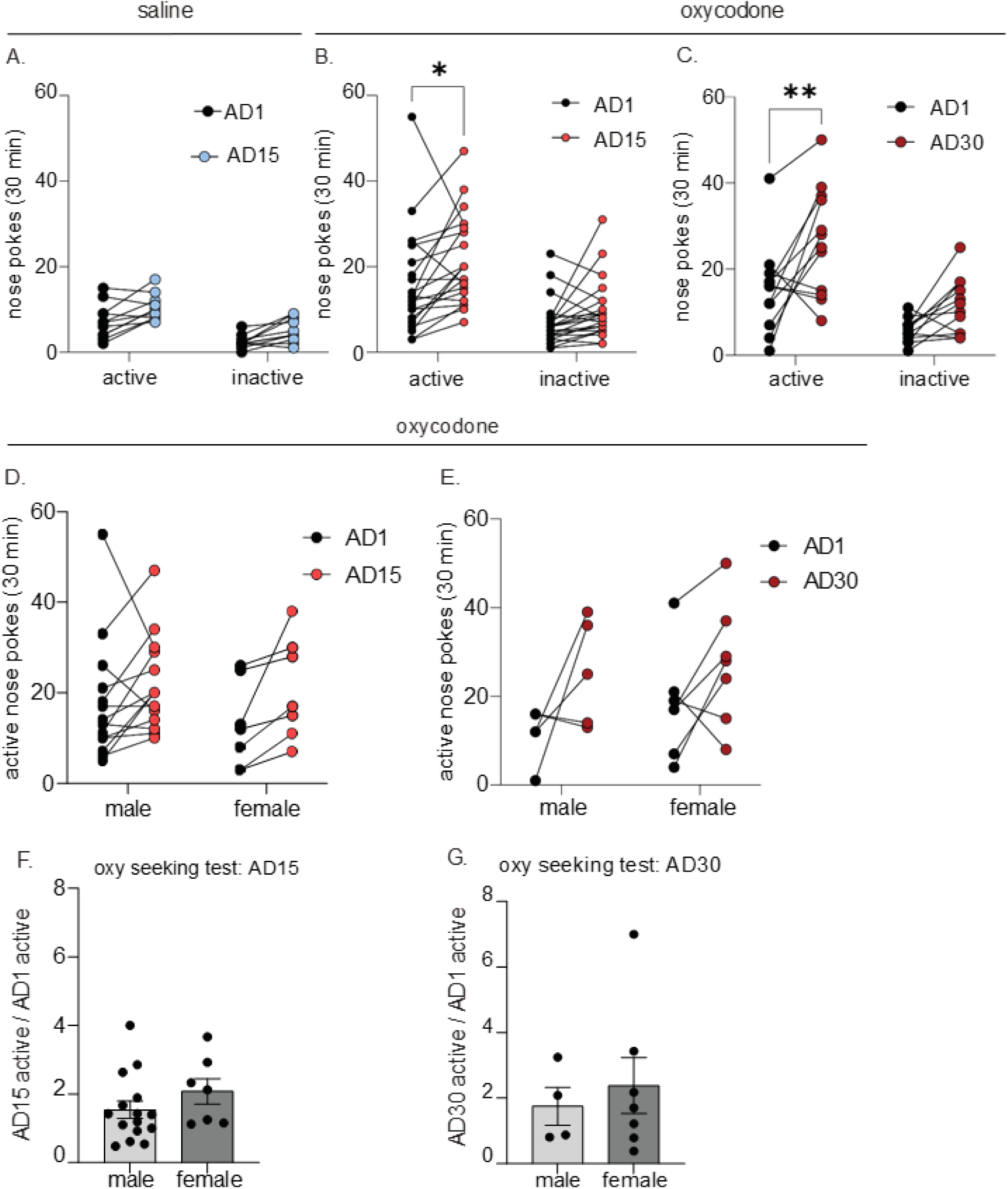
Incubation of oxycodone craving in male and female Long-Evans rats. A subset of rats from Fig. 1 received cue-induced seeking tests. (A) Saline self-administering animals did not show a time-dependent change in active hole responding between abstinence day (AD)1 and AD15 (*p*>0.05). (B) Oxycodone rats demonstrated enhanced oxycodone seeking on abstinence day (AD) 15 versus AD1 (active hole responding) but no time-dependent change in inactive hole responding (\**p*<0.05). (C) Enhanced oxycodone seeking persists on abstinence day 30 (\*\**p<*0.01). (D, E) Males and female oxycodone rats do not differ significantly in number of active hole or inactive hole nose pokes on AD1, AD15, or AD30 (p>0.05 male vs female on each AD). (F,G) The magnitude of incubation was calculated by dividing the number of active hole pokes during the late abstinence seeking test (AD15 or AD30) by the number of active hole pokes during the AD1 seeking test. Data are presented as mean ± SEM with individual data points shown. Male and female rats did not differ in the incubation score on AD15 or AD30 (*p*>0.05). Saline rats, *n=*15 (9 males, 6 females); AD1 and AD15 oxycodone rats, *n*=21 (14 males, 7 females); AD1 and AD30 oxycodone rats, *n*=12 (5 males, 7 females). More information about statistical analysis is presented in the Results section.

**Figure 3.**
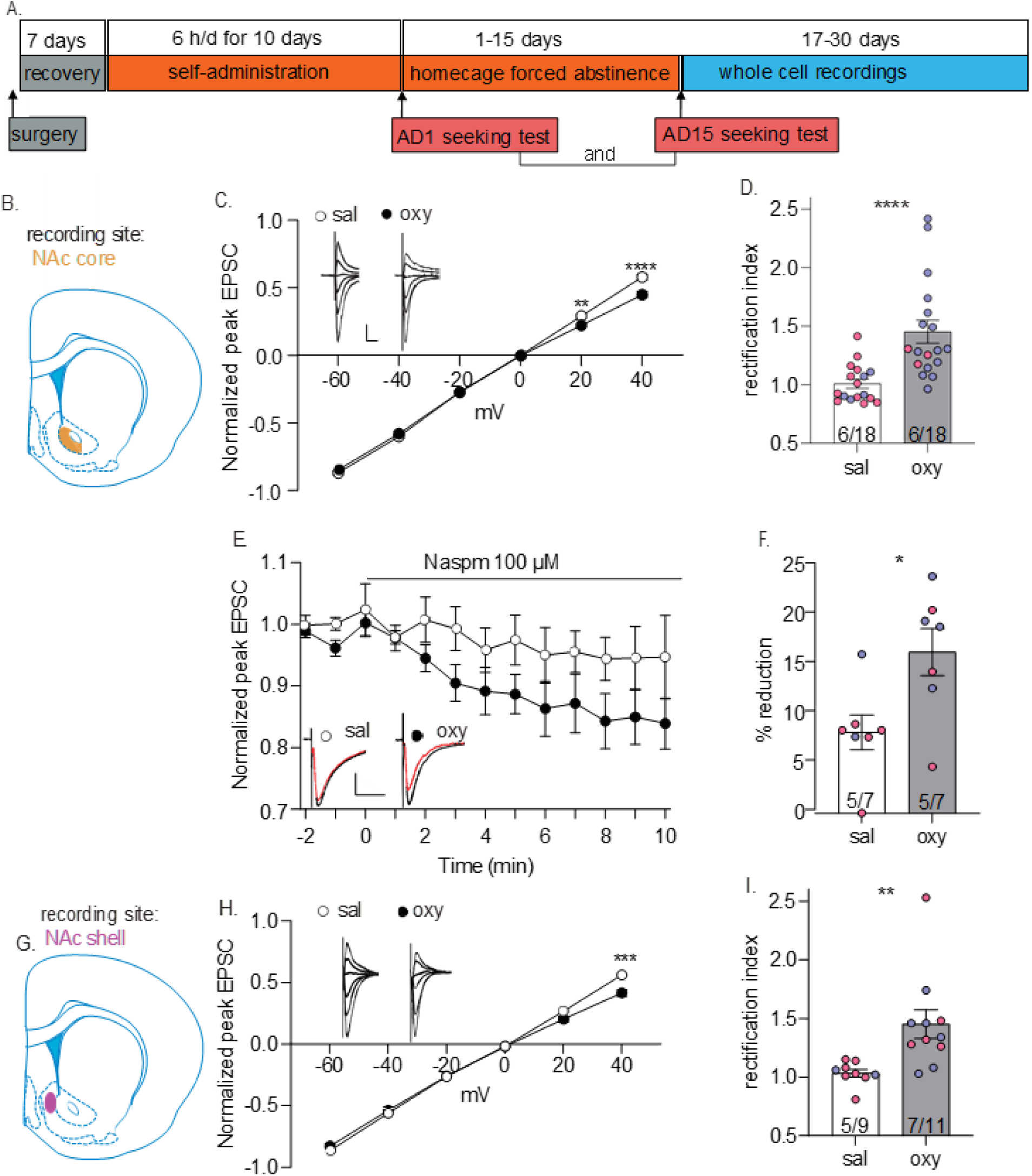
CP-AMPAR are upregulated in NAc core and shell synapses after incubation of oxycodone craving. (A) Experimental timeline. (B) Recording site for NAc core data in panels C-F. (C) I-V plot of AMPAR-mediated synaptic responses at different membrane holding potentials from NAc core MSNs on AD17-AD33 following saline (sal) or oxycodone (oxy) self-administration. The saline group showed a linear I-V relationship, while MSNs from oxy rats showed inwardly rectifying I-V relationships. Asterisks indicate significant reduction in EPSC amplitude relative to saline controls. Inset: example traces of AMPAR-mediated eEPSCs (top) at −60, −40, −20, 0, +20 and +40 mV. Scale bar: 100 pA/20 ms. (D) Quantification of rectification index (RI, [eEPSC-70mV/(−70-Erev)]/[eEPSC+40mV/(+40-Erev)]) showing a higher RI in NAc core MSNs from oxycodone rats compared to saline rats. Group sizes (also shown in bars as rats/cells): saline: 6 rats/18 cells; oxycodone: 6 rats/18 cells. (E) eEPSC amplitude (normalized to pre-Naspm baseline) during 100 μM Naspm application in NAc core MSNs from saline or oxycodone rats. Inset: representative eEPSC traces (black line: baseline; red line: 8-10 min after Naspm application). Scale bar: 100 pA/20 ms. (F) Mean eEPSC amplitudes measured 8-10 min after Naspm application. Naspm decreased eEPSC amplitudes in MSNs from oxycodone rats but not saline rats, indicating upregulation of CP-AMPARs in the former group. Group sizes (also shown in bars as rats/cells): saline: 5 rats/7 cells; oxycodone: 5 rats/7 cells. (G) Recording site for NAc shell data in panels H and I. (H) I-V plot of AMPAR-mediated synaptic responses of MSNs from NAc shell following 17-30 days of forced abstinence from saline or oxycodone self-administration. The saline group showed a linear I-V relationship, while MSNs from oxycodone rats showed inwardly rectifying I-V relationships. Asterisks indicate significant reduction in EPSC amplitude relative to saline controls. Inset: example traces of AMPAR-mediated eEPSCs. Scale bar: 100 pA/20 ms. (I) RI in NAc shell MSNs was significantly higher in oxycodone rats compared to saline rats. Group sizes (also shown in bars as rats/cells): saline: 5 rats/9 cells; oxycodone: 7 rats/11 cells. *p<0.05; **p<0.01; ***p<0.001; ****p<0.0001. Male (blue circles) and female (magenta circles) data were pooled for analysis. More information about statistical analysis is presented in the Results section.

### Tests for cue-induced oxycodone seeking

Oxycodone and saline rats received two cue-induced seeking tests: one test on AD1 (all rats) and a second test on either AD15 or AD30. For the seeking test, rats were returned to their self-administration chambers and were tested for 30 min or 1 hour under extinction conditions (i.e., nose pokes in the active hole resulted in presentations of the light cue previously paired with oxycodone, but oxycodone was not delivered). The number of responses in the active hole was used as a measure of drug seeking behavior. Responses in the inactive hole were also recorded but had no consequences. Rats that received AD1 and AD15 tests were used for electrophysiological studies between AD17 and AD33. Rats recorded on AD1-2 did not receive seeking tests.

### Electrophysiology

Whole-cell patch clamp recordings were performed in NAc medium spiny neurons (MSN) as in our prior work (e.g., (Conrad et al., 2008; McCutcheon et al., 2011b; McCutcheon et al., 2011a; Loweth et al., 2014; Scheyer et al., 2016; Kawa et al., 2022; Funke et al., 2023; Hwang et al., 2024; Wunsch et al., 2024)). AMPAR-mediated EPSCs were isolated pharmacologically. A bipolar tungsten electrode placed ∼300 µm from the recording site was used to evoke EPSCs. Only cells exhibiting a stable synaptic response (<15% variability) during 10-15 min of baseline recording were used. The contribution of CP-AMPARs to synaptic transmission was tested by determining IV relationships and calculating the rectification index (RI) for each cell: [EPSC_-70mV_/(−70 - E_rev_)]/[EPSC_+40mV_/(+40 - E_rev_)] (Kamboj et al., 1995). Both CP-AMPARs and CI-AMPARs contribute to EPSCs at −70mV holding potential (EPSC_-70mV_), but the contribution of CP-AMPARs is reduced at +40mV (EPSC_+40mV_) due to voltage-dependent block by intracellular polyamines. Thus, an increase in CP-AMPARs is reflected by a higher RI (see our prior work cited above). To confirm RI findings, we used the selective CP-AMPAR antagonist Naspm (100 µM). Naspm reduces the evoked AMPAR EPSC_-70mV_ by ∼5% in MSN from saline rats but elicits a larger reduction after CP-AMPARs increase, e.g., 25-30% after incubation of cocaine craving (Conrad et al., 2008; Purgianto et al., 2013; Loweth et al., 2014) and ∼20% after incubation of Meth craving (Funke et al., 2023). Thus, if Oxy abstinence elevates the RI, it should also increase Naspm sensitivity.

### Statistical analyses

Clampfit (v11, Molecular Devices), Excel (Microsoft), Prism10 (GraphPad) and Illustrator (Adobe) were used for data analysis and figures. All data were assessed for normality using Shapiro-Wilk tests. For comparison of two groups, Student’s t-tests (independent unless otherwise indicated) were used for normally distributed data and a Mann Whitney test was used for non-normally distributed data sets. ANOVAs were utilized for comparing multiple groups followed by Tukey’s or Bonferroni post hoc multiple comparisons (type of ANOVA specified within Results). For all analyses, significance was set at p<0.05. The data were presented as group mean ± S.E.M. Sample sizes were selected based on a priori power analysis (G-Power) supplemented to offset attrition typically experienced (e.g., due to failed catheters). In some cases, post hoc analyses were also performed to assess effect size for sex, and determine sample sizes that would be required in future studies of sex differences (Diester et al., 2019).

## Results

### Oxycodone self-administration and incubation of craving

Wildtype Long-Evans rats nose-poked to self-administer saline (15 rats; 9 male, 6 female) or oxycodone (58 rats; 36 male, 22 female) (6 h/d x 10 d; FR1; each infusion paired with a light cue) (Fig. 1A). We began by analyzing the number of nose pokes during self-administration using a mixed effects ANOVA with treatment (oxycodone or saline) as the between subjects factor, hole (active or inactive) as the within subjects factor, and session (1-10) as the repeated measure. The ANOVA revealed significant effects of session (F(9,648)=54.73, *p<*0.0001), treatment (F(1,72)=39.95, *p<*0.0001) and hole x session (F(9,609)=5.638, *p<*0.0001) and hole x session x treatment (F(9,609)=2.304, *p=*0.015) interactions. Oxycodone rats learned to discriminate between active and inactive holes and formed a strong preference for the oxycodone-associated hole, evidenced by significantly more active hole pokes compared to inactive pokes during all sessions (Bonferroni test, session 1 *p<*0.05, sessions 2-10 *p<*0.001; Fig. 1B). Saline rats also showed a significant preference for the active hole during sessions 1-4, 6, and 8 (Bonferroni test, *p*<0.05; Fig. 1B), although responding on the active hole was substantially lower in the saline group compared to the oxycodone group during sessions 7, 9 and 10 (Bonferroni test, *p*<0.05; Fig. 1B).

We compared nose pokes during oxycodone self-administration between males (*n =* 36) and females (*n =* 22) using a mixed model ANOVA with sex as the between subject factor, hole as the within subject factor, and session as the repeated measure. There was a main effect of session [F(9, 976)=2.637, *p=*0.0051] as well as hole [F(1,112)=50.28, *p<*0.0001] but not sex (*p*=0.6287) (Fig. 1C). No significant interactions were observed. A mixed model ANOVA was then used to assess oxycodone infusions during self-administration training with sex as the between subject factor and session as the repeated measure. This revealed a significant main effect of session [F(9,485)=4.073, *p<*0.0001] but not sex ([F(1,56)=0.05907, *p=*0.8089]. While male rats showed significantly more infusions on day 10 vs day 5 (Bonferroni test, *p*=0.037), females did not show a significant increase in infusions across sessions. We also compared average oxycodone infusions over the last 5 days of self-administration and found no significant difference between male and female rats (t-test, *p*=0.8324, Fig. 1E). Overall, these data suggest no pronounced sex differences in oxycodone self-administration although escalation of intake may be more pronounced in males.

A subset of the rats depicted in Fig. 1 underwent home cage forced abstinence and were tested for cue-induced oxycodone seeking (Fig. 2). Each of these rats received 2 cue-induced seeking tests, either on AD1 and AD15 (saline: 14 rats, 5 male, 9 female; oxycodone: 21 rats, 14 male, 7 female) or on AD1 and AD30 (oxycodone: 12 rats, 5 male, 7 female). The number of nose pokes during the seeking tests was analyzed with a mixed model ANOVA with treatment as the between subjects factor, hole as the within subjects factor, and abstinence day (AD1 or AD15) as the repeated measure. The ANOVA revealed main effects of abstinence day (F(1,50)=14.82, *p<*0.0003), hole (F(1,33)=14.40, *p*=0.0004), and treatment (F(1,50)=62.11, *p<*0.0001). Interactions were observed between hole x treatment (F(1,33)=10.72, *p*=0.0025), hole x abstinence day (F(1,44)=5.299, *p*=0.0261), and abstinence day x treatment (F(1,44)=7.120, *p*=0.011). Oxycodone rats expressed incubation of craving, demonstrated by significantly more active hole responses on AD15 relative to AD1 (*p*<0.05). Oxycodone rats also discriminated between the active and inactive ports, revealed by more active nose pokes than inactive pokes on AD1 (*p<*0.0001) and AD15 (*p<*0.0001) (Fig. 2B). By contrast, saline self-administering rats (*n=*11) did not discriminate between the active and inactive holes during either seeking test (*p>*0.05), nor did they significantly increase their active hole responding on AD15 versus AD1 (*p* >0.05) (Fig. 2A).

A two-way ANOVA for nose pokes was performed with hole as the within subjects factor and abstinence day (AD1 and AD30) as the repeated measure. Treatment was not included as factor because no saline rats were tested on AD30. We observed a main effect of abstinence day (F(1, 44)=11.04, *p=*0.0018) and hole (F(1,44)=23.62, *p<*0.0001) with active hole responses significantly increased on AD30 relative to AD1 (*p=*0.0063) (Fig. 2C) confirming that incubation of oxycodone craving persists to at least AD30. Further analysis showed that the magnitude of incubation for individual rats did not show a significant correlation with oxycodone infusions during the last 5 days of self-administration (Fig. S2).

To assess sex as a biological variable (SABV), we compared active hole responses between male and female oxycodone rats during the AD1, AD15 and AD30 seeking tests using a two-way ANOVA with sex and abstinence day as between subjects factors. Results showed a main effect of abstinence day (F(1,48)=3.301, *p=*0.0454) but not sex (*p>*0.05) There were no significant difference in the incubation score between males and females on either AD15 (Fig. 2F) or AD30 (Fig. 2G) (t-tests, *p*>0.05), in agreement with prior studies on incubation of opioid craving (see Discussion).

### CP-AMPAR upregulation in NAc core and shell after incubation of Oxy seeking

To determine if CP-AMPAR accumulate in NAc synapses alongside incubation, we performed whole cell recordings from a subset of the saline and oxycodone rats shown in Fig. 2, targeting the AD15-30 period when incubation is stably expressed. The timeline for electrophysiological experiments is shown in Fig. 3A. For core, we recorded around the anterior commissure as in our studies after cocaine incubation (Fig. 3B) (Conrad et al., 2008; McCutcheon et al., 2011a; Hwang et al., 2024). The shell is a heterogeneous structure (Al-Hasani et al., 2015; Berridge and Kringelbach, 2015; Castro and Bruchas, 2019); we focused on medial shell based on prior studies of glutamate transmission in opioid reward and aversion (Fig. 3G) (Zhu et al., 2016; Madayag et al., 2019).

Beginning with MSN in the NAc core, we generated I-V curves for the AMPAR-mediated evoked EPSC (eEPSC) to assess inward rectification, a hallmark of CP-AMPARs. MSNs from saline rats exhibited a linear I-V relationship, whereas the oxycodone group showed inward rectification (Fig. 3C). A two-way ANOVA with treatment (saline or oxycodone) as a between-subject factor and voltage (−60, −40, −20, 0, +20, +40 mV) as a within-subject factor showed a main effect of treatment (F(1,204)=10.53, *p*=0.0014) and voltage (F(5,204)=2725, *p*<0.0001), and a treatment x voltage interaction (F(5,204)=9.310, *p*<0.0001). The normalized eEPSC was significantly reduced in MSNs from oxycodone rats at +20 mV (*p*=0.005) and +40 mV (*p*<0.0001). Consistent with these findings, the oxycodone group showed a significantly elevated rectification index (unpaired *t*-test, *t*(33)=4.075, *p*=0.0003) (Fig. 3D).

We extended our investigation of CP-AMPAR to the NAc shell. As found for core, we detected significant inward rectification in NAc shell MSNs from oxycodone rats, but not saline rats (unpaired *t*-test, *t*(18)=3.033, *p*=0.0076) (Fig. 3H, I). A two-way ANOVA revealed a main effect of voltage (F(5,90)=1099, *p*<0.0001) and a treatment x voltage interaction (F(5,90)=5.222, *p*= 0.0003) with the normalized eEPSC at +40 mV in the oxycodone group significantly lower than the saline group (Bonferroni, *p*<0.0001).

To confirm the presence of CP-AMPAR, eEPSCs from NAc core MSNs were measured before and after bath application of the selective CP-AMPAR antagonist Naspm (100 μM) (Fig. 3E,F). Mean eEPSC amplitude, measured 10 min after Naspm application and expressed as percent of baseline, was significantly reduced in the oxycodone group compared to saline controls (unpaired *t*-test, *t*(12)=2.734, *p*=0.0181).

In some of the rats used for assessment of inward rectification or Naspm sensitivity, we also assessed the paired pulse ratio during baseline recordings. Saline and oxycodone groups showed no difference in paired pulse ratio in either core (saline: 1.0891±0.0717, 13 cells/5 rats; oxycodone: 1.0648±0.0826, 29 cells/12 rats) or shell (saline: 1.05±0.0781, 8 cells/4 rats; oxycodone: 1.01±0.0597, 9 cells/5 rats (sexes combined for PPR data; unpaired t-tests for each subregion *p*>0.05), indicating that the postsynaptic adaptations shown in Fig. 3 are not accompanied by changes in release probability.

Since we typically record one rat per day, recording over a range of AD’s has a practical advantage in enabling us to run a cohort of rats through self-administration and have a reasonable period of time to do recordings. However, although we showed that incubation is stable from AD15 to at least AD30 (Fig. 2), it was necessary to establish whether plasticity is also stable. To address this, we tested the relationship between the AD on which the rat was recorded (range was AD17-33) and the average RI of cells recorded from the rat. We did not observe a significant correlation (Fig. S3A). Thus, CP-AMPAR upregulation, like oxycodone incubation (Fig. 2), is stable over the abstinence period targeted in our study. A second concern is that a series of oxycodone incubation studies, albeit using a longer duration self-administration regimen than ours, divided rats into low and high oxycodone-taking subgroups; these subgroups showed similar incubation but some gene expression differences in dorsal striatum and hippocampus (Blackwood et al., 2019b; Blackwood et al., 2019a; Salisbury et al., 2020). We therefore tested correlations between oxycodone intake (average infusions, last 5 d of self-administration) and average RI of cells recorded from each rat. We found no significant correlation between oxycodone intake and the average RI (Fig. S3B), indicating that CP-AMPAR upregulation occurs across a range of oxycodone intake levels.

In summary, electrophysiological results in unidentified MSN demonstrate upregulation of CP-AMPARs in core and shell subregions of the NAc in association with incubation of oxycodone craving and further establish that the magnitude of this upregulation is stable across AD17-33.

### Oxycodone self-administration and incubation in transgenic rats

To visualize D1-positive or A2a-positive MSN for slice recordings, we crossed D1-Cre and A2a-Cre rats (Pettibone et al., 2019) with ZsGreen or TdTomato reporter lines (denoted here as “reporter”), as in previous studies (Hwang et al., 2024). Prior to recordings, rats self-administered saline or oxycodone (6 h/d for 10 d). Using a two-way RM ANOVA with Bonferroni’s multiple comparisons test, we analyzed active and inactive nose-pokes for both D1-Cre/reporter rats [Fig. 4A; hole effect F(1,30)=24.90, *p*<0.0001; session effect F(2.958,142)=10.19, *p*<0.0001; treatment effect F(1,24)=6.949, *p*=0.0145; hole x session x treatment effect F(27,432)=3.466, *p*<0.0001] and for A2a-Cre/reporter rats [Fig. 4B; hole effect F(1,26)=33.87, *p*<0.0001; session effect F(3.818,190.9)=23.67, *p*<0.0001; treatment effect F(1,25)=4.179, *p*=0.0516; hole x session x treatment effect F(27,450)=4.821, *p*<0.0001]. Both D1-Cre/reporter and A2a-Cre/reporter rats that self-administered oxycodone discriminated between active and inactive ports during self-administration sessions 5-10 and sessions 4-10, respectively [Fig. 4A (D1-Cre/reporter): *p*<0.05 for session 10; *p*<0.01 for sessions 5, 7, and 9; *p*<0.001 for sessions 6 and 8; Fig 4B (A2a-Cre/reporter): *p*<0.05 for sessions 4 and 10; *p*<0.01 for session 5; *p*<0.001 for sessions 6, 8, and 9; *p*<0.0001 for session 7]. While the saline controls in both transgenic lines discriminated between the active and inactive hole [Fig. 4A (D1-Cre/reporter): *p*<0.0001 in session 1; Fig 4B (A2a-Cre/reporter): *p*<0.05 in sessions 1 and 4-8], the D1-Cre/reporter oxycodone rats exhibited significantly higher active nose pokes than their saline controls in sessions 5-10 (Fig. 4A: *p*<0.05 on sessions 6-8; *p*<0.01 in sessions 5, 9, and 10), while the A2a-Cre/reporter oxycodone rats exhibited significantly higher active nose pokes for oxycodone compared to their saline controls in sessions 6-9 (Fig. 4B: *p*<0.05 in sessions 6 and 9; *p*<0.01 in sessions 7 and 8). Furthermore, both D1-Cre/reporter and A2a-Cre/reporter rats showed significantly greater total infusions throughout the self-administration sessions compared to their respective saline controls [Fig. 4D (D1): treatment effect F(1,24)=14.62, *p*<0.001; session effect F(2.639,63.34)=5.806, *p*<0.01; treatment x session interaction F(9,216)=6.053, *p*<0.0001; Fig 4E (A2a): treatment effect F(1,25)=4.924, *p*<0.05; session effect F(2.436,60.90)=15.30, *p*<0.0001, treatment x session interaction F(9,225)=11.39, *p*<0.0001]. Importantly, we also compared D1-Cre/reporter and A2a-Cre/reporter lines. A two-way RM ANOVA revealed that the two lines showed no significant difference in the number of active nose pokes [Fig. 4C: transgenic line effect F(1,28)=0.7664, *p*>0.05] nor the number of infusions [Fig 4F: transgenic line effect F(1,28)=1.987, *p*>0.05].

**Figure 4.**
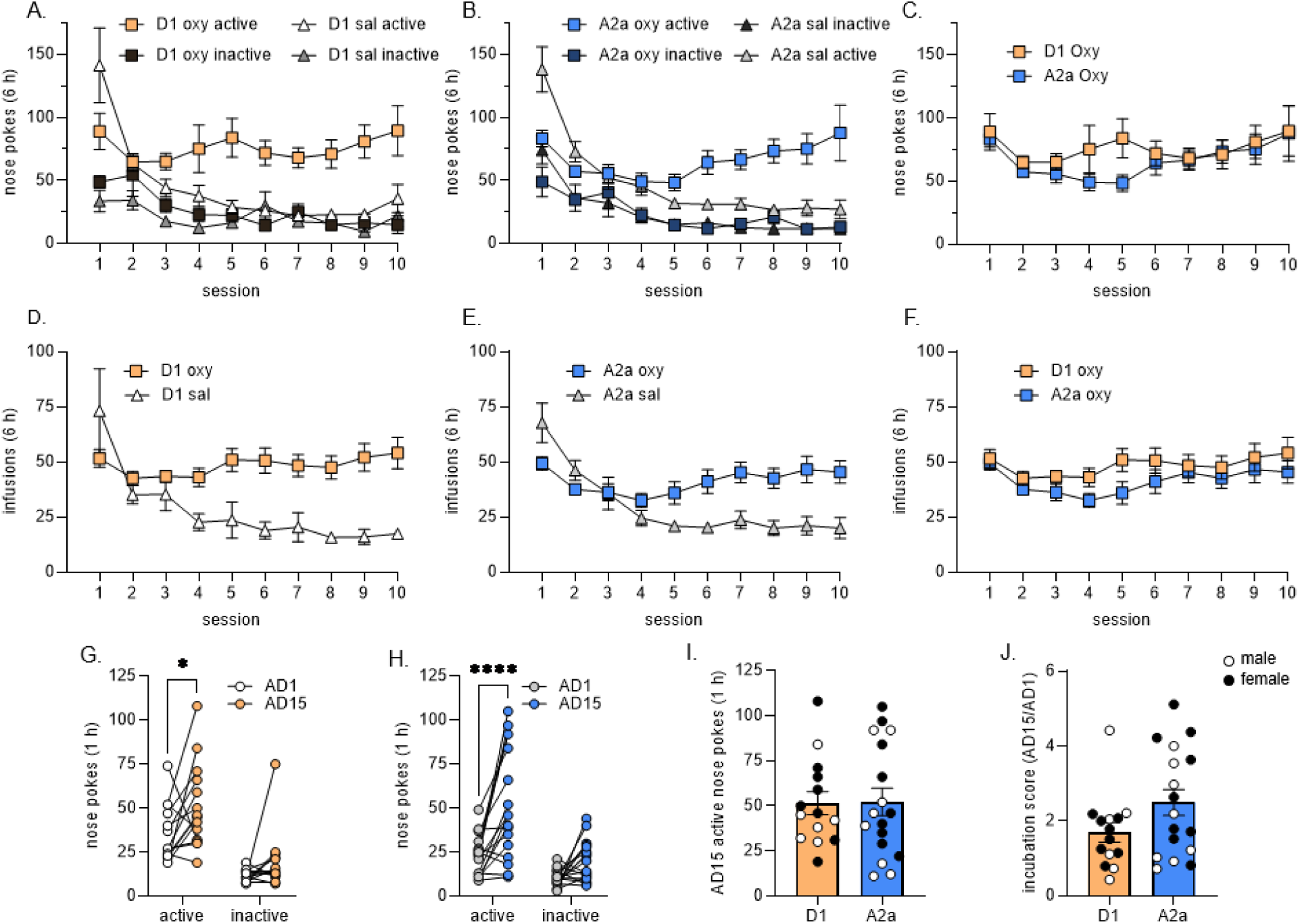
Oxycodone self-administration and incubation of craving in transgenic rat lines. D1-Cre or A2a-Cre rats were crossed with ZsGreen or TdTomato reporter lines. Experiments were performed in Cre+ offspring. Rats were trained to self-administer oxycodone or saline and reach rat received two cue-induced seeking tests [abstinence day (AD)1 and AD15], as shown for WT rats in the Fig. 1A timeline. (A) Active and inactive nose pokes for D1-Cre/reporter rats self-administering saline or oxycodone (active vs inactive pokes for oxycodone rats, *p<*0.05 on sessions 5-10). (B) Active and inactive nose pokes for A2a-Cre/reporter rats self-administering saline or oxycodone (active vs inactive pokes for oxycodone rats, *p<*0.05 on sessions 4-10). (C) Active nose pokes did not differ significantly between oxycodone self-administering D1-Cre/reporter rats and A2a-Cre/reporter rats (p>0.05). (D) Number of infusions across all self-administration sessions for D1-Cre/reporter rats (oxycodone > saline infusions, p<0.05 for sessions 4-10). (E) Number of infusions across all self-administration sessions for A2a-Cre/reporter rats (oxycodone > saline infusions, p<0.05 for sessions 5-10). (F) Oxycodone infusions during self-administration training did not differ between the two transgenic rat groups (*p*>0.05). Data for panels A-F are presented as mean ± SEM. (G, H) Oxycodone self-administering rats in both D1-Cre/reporter (G) and A2a-Cre/reporter (H) transgenic lines demonstrated higher oxycodone seeking on AD15 versus AD1 (active hole responding) but no time-dependent change in inactive hole responding [(G) *t*(26)=2.681, \**p*<0.05; (H) *t*(32)=5.435, \*\*\*\**p*<0.0001 vs AD1, paired t-tests]. (I) Active hole responding on AD15 did not differ between the two transgenic lines (*p*>0.05, t-test). (J) Incubation score (see Fig. 1 for definition) did not differ between the two transgenic lines (*p*>0.05, t-test). Males and females indicated by white and black circles respectively. D1 saline rats, *n* = 10 (5 males, 5 females), D1 oxycodone rats, *n* = 14 (7 males, 7 females), A2a saline rats, *n* = 13 (5 males, 8 females), A2a oxycodone rats, *n* = 17 (8 males, 9 females). More information about statistical analysis is presented in the Results section.

All rats received 60 min cue-induced seeking tests on AD1 and on AD15 to assess incubation. Assessment of nose pokes in the previously active hole using a two-way RM ANOVA with Sidak’s multiple comparison’s test revealed incubation of oxycodone craving in D1-Cre/reporter and A2a-Cre/reporter lines [Fig. 4G (D1-Cre/reporter): hole effect F(1,26)=34.90, *p*<0.0001; AD effect F(1,26)=7.970, *p*<0.01; hole x AD interaction F(1,26)=0.9378, *p*>0.05; Fig. 4H (A2a-Cre/reporter): hole effect F(1,32)=21.78, *p*<0.0001; AD effect F(1,32)=26.79, *p*<0.0001; hole x AD interaction F(1,32)=6.30, *p*<0.05]. There was no significant difference in AD15 responding between the two rat lines (Fig. 4I, Mann-Whitney test, U=115 *p*>0.05). Furthermore, there was no significant difference in the magnitude of incubation as reported by the incubation score (AD15/AD1) between the D1-Cre/reporter and A2a-Cre/reporter rats (Fig. 4J, Mann-Whitney test, U=84.5 *p*>0.05).

Finally, comparison of male and female oxycodone rats within the D1-Cre/reporter group using a two-way RM ANOVA revealed no significant differences in active hole responding [F(1,14)=3.332, *p*>0.05], inactive hole responding [F(1,14)=3.864, *p*>0.05] or infusions [F(1,14)=1.157, *p*>0.05] during self-administration training; the same held for their saline controls [active hole F(1,8)=0.1207, *p*>0.05; inactive hole F(1,8)=2.074, *p*>0.05, infusions F(1,8)=0.016, *p*>0.05]. Using the same analysis, no sex differences for active hole responding [F(1,12)=0.1230, *p*>0.05], inactive hole responding [F(1,12)=0.7954, *p*>0.05], or infusions [F(1,12)=0.2982, *p*>0.05] were found for oxycodone rats in the A2a-Cre/reporter group; the same held for their saline controls [active hole F(1,11)=0.005, *p*>0.05; inactive hole F(1,11)=1.214, *p*>0.05; infusions F(1,11)=0.043, *p*>0.05]. These data demonstrate robust oxycodone self-administration and incubation of craving in males and females from our transgenic rat lines.

### CP-AMPARs upregulate in both D1 and A2a/D2 MSN

A subset of rats from Fig. 4 were recorded to assess AMPAR transmission in D1+ MSN (fluorescent cells from D1-Cre/reporter rats) and A2a+ MSN (fluorescent cells from A2a-Cre/reporter rats) on AD17-33 following saline or oxycodone self-administration. For a representative image of a patched D1+ MSN, see (Hwang et al., 2024). Using a two-way RM ANOVA to evaluate the MSN in the NAc core, we observed linear I-V relationships in both MSN subtypes for saline rats and significant inward rectification in both D1+ MSN and A2a+ MSN after oxycodone incubation [Fig. 5A (D1+ MSN): voltage effect F(2.801,106.5)=2420, *p*<0.0001; treatment effect F(1,38)=11.75, *p*<0.01; voltage x treatment interaction F(5,190)=3.815, *p*<0.01; Fig. 5B (A2a+ MSN): voltage effect F(3.076,116.9)=4464, *p*<0.0001; treatment effect F(1,38)=6.638, *p*<0.05; voltage x treatment interaction F(5,190)=21.62, *p*<0.0001]. We also assessed AMPAR transmission in the NAc shell and observed linear I-V relationships for saline controls but inward rectification for both MSN subtypes after oxycodone incubation [Fig. 5C (D1+ MSN): voltage effect F(2.508,57.68)=1512, *p*<0.0001; treatment effect F(1,23)=3.938, *p*=0.0593; voltage x treatment interaction F(5,115)=4.320, *p*<0.01; Fig. 5D (A2a+ MSN): voltage effect F(2.270, 52.21)=1576, *p*<0.0001; treatment effect F(1,23)=12.48, *p*<0.01; voltage x treatment interaction F(5,115)=5.315, *p*<0.001]. These findings are reflected by an elevated RI for D1+ MSN and A2a+ MSN after oxycodone incubation relative to respective saline controls in both subregions [Fig. 5E (Core): One-way ANOVA, F(3,84)=12.94, *p*<0.0001; Fig. 5F (Shell): One-way ANOVA, F(3,48)=8.202, *p*<0.001].

**Figure 5.**
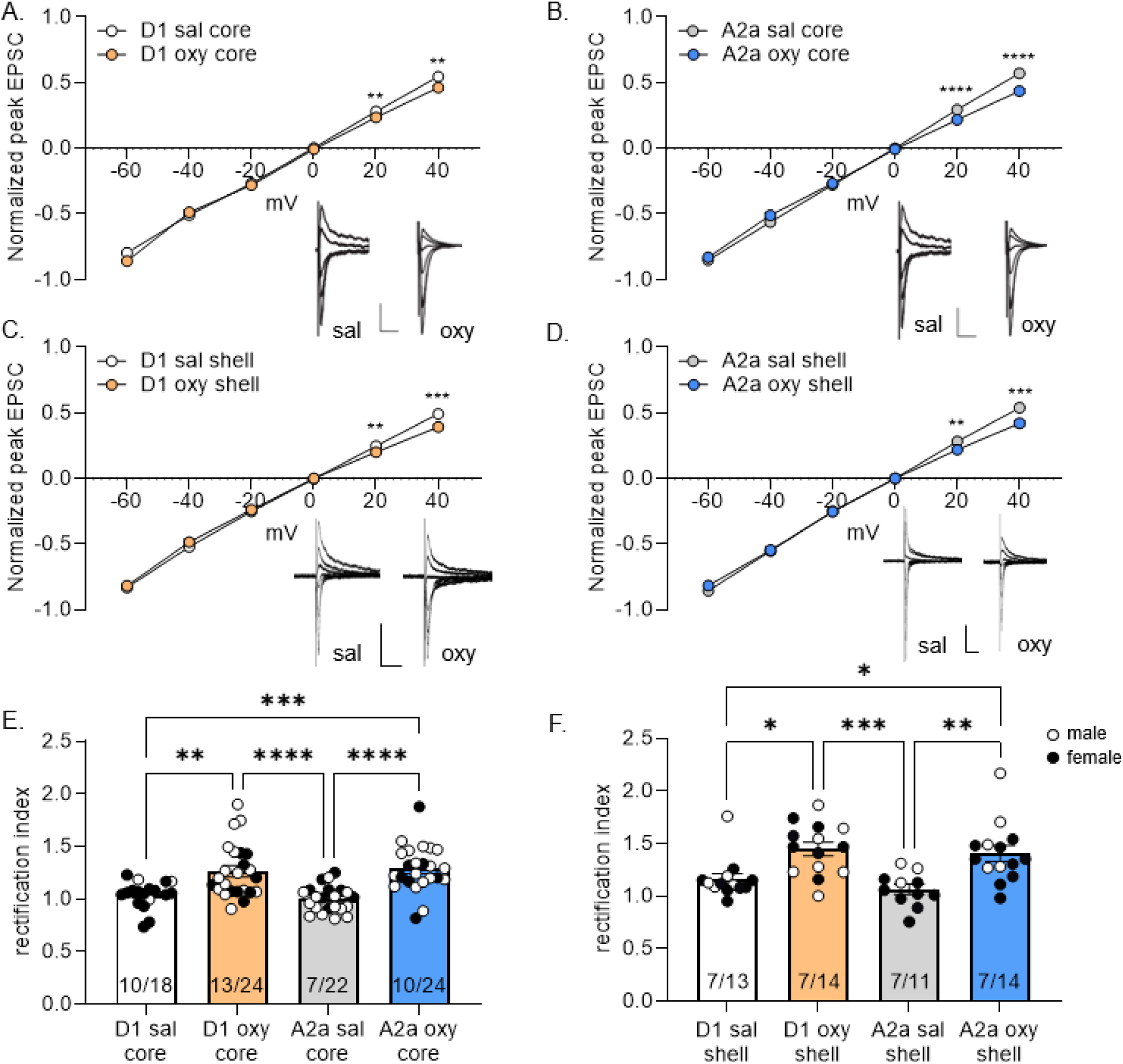
CP-AMPARs are upregulated in D1 and A2a medium spiny neurons (MSN) in NAc core and shell synapses after incubation of oxycodone craving. A subset of D1-Cre/reporter and A2a-Cre/reporter rats from Figure 4 was used for these studies. Whole-cell patch clamp recordings were performed in core and shell MSN on AD17-AD33 following extended-access saline or oxycodone self-administration as shown in the Fig. 3A timeline. (A,B) I-V plots of AMPAR-mediated synaptic responses at different membrane holding potentials from D1+ MSN (A) and A2a+ MSN (B) in NAc core. Saline groups showed a linear I-V relationship, while both D1 and A2a MSN from oxycodone rats showed inwardly rectifying I-V relationships. Asterisks indicate significant reduction in EPSC amplitude relative to saline controls. Inset: example traces of AMPAR-mediated eEPSCs at −60, −40, −20, 0, +20 and +40 mV. Scale bar: 100 pA/20 ms. Group sizes (also shown in bars as rats/cells): D1 Saline: 10 rats/18 cells; D1 oxycodone: 13 rats/24 cells; A2a saline: 7 rats/22 cells; A2a oxycodone: 10 rats/24 cells. (C,D) I-V plots of AMPAR-mediated synaptic responses from D1+ MSN (C) and A2a+ MSN (D) in NAc shell. Saline groups showed a linear I-V relationship, while both D1+ and A2a+ MSN from oxycodone rats showed inwardly rectifying I-V relationships (significant differences from saline indicated by asterisks). Inset and scale bar are the same as described above. Group sizes (also shown in bars as rats/cells): D1 Saline: 7 rats/13 cells; D1 oxycodone: 7 rats/14 cells; A2a saline: 7 rats/11 cells; A2a oxycodone: 7 rats/14 cells. (E,F) Quantification of the rectification index (RI) demonstrated that all oxycodone groups (D1+ and A2a+ MSN in both core and shell) had significantly higher RI values than their respective saline controls. *p<0.05; **p<0.01; ***p<0.001; ****p<0.0001. Male (white circles) and female (black circles) data were pooled for analysis. More information about statistical analysis is presented in the Results section.

We also performed recordings on AD1-2 to confirm that the CP-AMPAR upregulation demonstrated in Fig. 5 requires a period of forced abstinence to become apparent, as would be expected from the ability of intra-NAc core or shell injection of the CP-AMPAR antagonist Naspm to inhibit incubated cue-induced oxycodone seeking (AD15 or AD30) without affecting cue-induced seeking on AD1 (Wong et al., 2022) (see Introduction). Preliminary data indicate that the RI of both MSN subtypes in NAc core on AD1-2 was at saline control levels (∼1; see Fig. 3) based on recordings in A2a+ (fluorescent) and A2a-(non-fluorescent; presumed D1) MSN from the NAc core of A2a-Cre/TdTomato rats (A2a+: RI = 1.09 ± 0.03, 9 cells/2 rats; A2a-: RI = 1.04 ± 0.03, 7 cells/2 rats). Although we are presently collecting more AD1-2 data, these results are consistent with time-dependent CP-AMPAR accumulation in both MSN subtypes during Oxy incubation.

Our primary statistical analysis for Fig. 5 focused on the effect of treatment, using pooled male/female data. However, for core A2a+ MSN (where both sexes are well represented in saline and oxycodone groups), secondary analyses were done to explore potential sex differences, using a published approach (Diester et al., 2019). First, we analyzed RI data by ANOVA with treatment and sex as factors. No significant main effects were found. However, effect size analysis confirmed that we were well powered to detect an effect of treatment (oxycodone versus saline) [effect size (f) 0.733, power 0.99]. For sex, however, the effect size was small (f=0.148); accordingly, calculations indicated that ∼180 cells/sex would be needed to achieve a power of 0.8. Thus, at most, there may be very small sex differences in CP-AMPAR levels in core A2a MSN of Oxy vs. Sal rats. Future studies could explore this for both NAc subregions and additional AD periods. In prior studies, our results did not suggest sex differences in CP-AMPAR upregulation in unidentified NAc core MSN after incubation of cocaine or methamphetamine craving (Kawa et al., 2022; Funke et al., 2023; Hwang et al., 2024) or in D1 or A2a MSN after cocaine incubation using the same transgenic lines employed here (Hwang et al., 2024).

## DISCUSSION

Here we show for the first time that CP-AMPARs accumulate in excitatory synapses on both D1 and A2a/D2 MSN in both core and shell NAc subregions during oxycodone incubation, i.e., CP-AMPAR levels are low on AD1-2 but stably elevated on AD17-33. These findings explain the ability of the CP-AMPAR antagonist Naspm, infused into NAc core or shell, to prevent expression of oxycodone incubation (Wong et al., 2022). This contrasts with psychostimulant incubation, where CP-AMPARs upregulate in both NAc subregions but nearly all evidence suggests selectivity for D1 MSN (see below). Aside from our work [(Wong et al., 2022) & present findings], there have been only a few studies on the NAc’s role during abstinence from Oxy self-administration. One implicated the NAc in oxycodone seeking during abstinence from oxycodone self-administration although incubation was not demonstrated (Gobin et al., 2022), while another found that incubation was associated with synaptic plasticity at paraventricular nucleus of the thalamus (PVT) inputs to NAc shell MSN (Alonso-Caraballo et al., 2024) (see below for more discussion). Finally, one study examined immediate early gene and growth factor gene expression changes associated with incubation (Blackwood et al., 2019a). The NAc has been studied after heroin (Kuntz et al., 2008; Airavaara et al., 2011; Theberge et al., 2012; Roura-Martinez et al., 2020; Xu et al., 2021) and morphine (Gillespie et al., 2022; Mayberry et al., 2022a) incubation but from the standpoint of gene expression not synaptic function. However, glutamatergic plasticity including CP-AMPAR upregulation in the NAc has been implicated in other opioid addiction models (see next section).

### Prior work implicates plasticity in D1 and A2a MSN in opioid reward and aversion

Our understanding of D1 and A2a MSN outputs and their activity in behavioral states has evolved in recent years, leading to updating of the canonical model of a direct pathway (D1) that promotes action and an indirect pathway (A2a) that opposes it. Current models emphasize the necessity for integrated D1 and A2a MSN activity in shaping motivated behavior (Burke et al., 2017; Kupchik and Kalivas, 2017; Bariselli et al., 2019; Fox and Lobo, 2019; Allichon et al., 2021; Zachry et al., 2023; Fang and Creed, 2024). Nonetheless, the prevailing view is that plasticity tipping the balance of excitatory drive towards D1 MSN promotes motivated behavior while relatively greater drive to A2a MSN is associated with aversion. Results after different opioid regimens generally support this, including studies of MSN using fiber photometry (O’Neal et al., 2022), chemogenetic or optogenetic manipulations (O’Neal et al., 2020; Giannotti et al., 2021), Fos (Enoksson et al., 2012), cAMP signaling (Muntean et al., 2019), and electrophysiology - for the latter, relatively greater excitatory drive to NAc shell D1 MSN was found after repeated i.p. morphine injections leading to sensitization or CPP (Graziane et al., 2016; Hearing et al., 2016; Madayag et al., 2019), while regimens leading to dependence and withdrawal elicited a shift towards greater excitatory drive to A2a MSN (Zhu et al., 2016; Madayag et al., 2019; Zhu et al., 2023). However, this pattern of D1/reward and A2a/aversion is not absolute, e.g., increased excitatory drive to core D1 MSN contributes to negative affective states after 5 d of fentanyl drinking (Fox et al., 2022).

Our results are the first to link upregulation of CP-AMPARs in the NAc to the incubation of cue-induced opioid seeking during abstinence from opioid self-administration. We were motivated to explore their potential role for two reasons: 1) strong evidence for CP-AMPAR mediation of psychostimulant incubation (see next section), and 2) prior reports of opioid-induced CP-AMPAR upregulation in the NAc shell. In the latter studies, the MSN subtype showing CP-AMPAR plasticity depended on the opioid regimen. After i.p. morphine injections leading to sensitization or CPP, excitatory drive to shell D1 MSN is strengthened in part by CP-AMPAR insertion, but this is not observed in shell A2a MSN, or core D1 or A2a MSN (Hearing et al., 2016). Likewise, CP-AMPARs are recruited to NAc shell synapses after fentanyl CPP followed by extinction training (Panopoulou and Schluter, 2022). In the only opioid self-administration study to assess CP-AMPARs, their levels increased in PVT-shell D1 but not A2a synapses after heroin self-administration and extinction training (Paniccia et al., 2024). In contrast, after escalating dose regimens used to elicit opioid withdrawal-associated aversive states (such regimens deliver higher opioid doses compared to sensitization, CPP or self-administration regimens), CP-AMPARs increase in A2a but not D1 MSN (Zhu et al., 2016). Specifically, CP-AMPARs increased in PVT-shell A2a MSN (but not D1 MSN) synapses and this plasticity was necessary for expression of aversive effects of morphine withdrawal (Zhu et al., 2016). NAc shell CP-AMPARs were also implicated in aversive effects of morphine withdrawal using biochemical and behavioral methods (Russell et al., 2016). Another electrophysiological study after withdrawal from an escalating dose morphine regimen found increased mEPSC frequency but not amplitude in shell A2a MSN, with no changes in shell D1, core D1, or core A2a MSN (Madayag et al., 2019).

The use of different opioids and regimens in these studies complicates their integration. Overall, however, they suggest that sensitization/CPP/self-administration regimens can be associated with elevated CP-AMPARs in shell D1 MSN, while aversive states of opioid withdrawal can be associated with elevated CP-AMPARs in shell A2a MSN. It is possible that our results can be interpreted in this framework, with CP-AMPAR elevation in D1 MSN driving incubation and CP-AMPAR elevation in A2a MSN underpinning negative affective states that may persist during forced abstinence from oxycodone self-administration. Experiments are underway to evaluate this possibility. It is also interesting that CP-AMPAR plasticity occurs in core and shell after oxycodone self-administration (present results) but preferentially in shell after non-contingent opioid administration (see above). As suggested previously (Hearing, 2019), it makes sense that the core would be more engaged in studies in which the opioid is self-administered given the core’s role in goal-directed behavior.

### CP-AMPAR plasticity in psychostimulant versus opioid incubation

Although incubation of craving occurs across drug classes, the incubation of cocaine craving has been best studied. It is well established that CP-AMPARs accumulate in MSN synapses in both core and shell subregions during forced abstinence from cocaine self-administration and that expression of cocaine incubation depends on CP-AMPAR activation in both subregions (Conrad et al., 2008; Lee et al., 2013; Loweth et al., 2014; Ma et al., 2014; Ma et al., 2016; Kawa et al., 2022; Hwang et al., 2024). The requirement for CP-AMPAR upregulation in both core and shell may relate to spiraling loops connecting them (Haber, 2016) or perhaps direct connections (van Dongen et al., 2008). Interestingly, the mechanisms leading to CP-AMPAR upregulation during cocaine incubation differ between the two subregions. Although it should be noted that direct comparison for some key endpoints are lacking (Wang et al., 2018; Christian et al., 2021; He et al., 2023), considerable evidence indicates that CP-AMPAR upregulation in the core involves retinoic acid-dependent homeostatic plasticity triggered by the contrast between high activity in NAc circuits during drug self-administration and lower activity during abstinence (Hwang et al., 2024; Wunsch et al., 2024) whereas in the shell cocaine self-administration generates silent synapses which in some pathways mature via CP-AMPAR insertion (Wright and Dong, 2020; Zinsmaier et al., 2022) (for review of subregion differences, see (Wolf, 2025)). Interestingly, silent synapses in NAc shell may be generated through different mechanisms after cocaine versus morphine administration (Graziane et al., 2016). It will be important to investigate mechanisms underlying CP-AMPAR upregulation in both subregions during incubation of Oxy craving. It will also be important to explore differences in subregion and MSN subtype plasticity between drug classes. Thus, the present results show that oxycodone incubation is associated with CP-AMPAR upregulation in both D1 and A2a MSN of core and shell, whereas it occurs selectively in D1 MSN of NAc core after cocaine incubation (Hwang et al., 2024) and methamphetamine incubation (E.-K. Hwang, M.M. Beutler, M.E. Wolf, Society for Neuroscience abstract 2024). Selective synaptic strengthening in D1 MSN after cocaine self-administration and abstinence was also observed in NAc shell (Pascoli et al., 2014) although more complex results were obtained after an extremely high dose cocaine regimen (Terrier et al., 2015).

Finally, it is important to compare pathway-specificity of incubation-related NAc plasticity across drug classes. Interestingly, input pathways showing CP-AMPAR upregulation differ for cocaine incubation (Lee et al., 2013; Ma et al., 2014; Pascoli et al., 2014; Terrier et al., 2015; Neumann et al., 2016) versus methamphetamine incubation (E.-K. Hwang, M.M. Beutler, M.E. Wolf, Society for Neuroscience abstract 2024). In a recent study of synapses between PVT inputs and unidentified NAc shell MSN after oxycodone self-administration, no changes in excitatory synaptic transmission were observed on AD1 (Alonso-Caraballo et al., 2024). However, after incubation of oxycodone craving (AD14), EPSC amplitude (input-output) and glutamate release probability were increased, but the AMPA/NMDA ratio and the rectification index were unchanged, indicating no CP-AMPAR upregulation at PVT-shell synapses. Increased NAc shell MSN intrinsic excitability was also observed on AD14 but not AD1 (Alonso-Caraballo et al., 2024). In contrast, our preliminary results indicate CP-AMPAR upregulation in PVT-shell (but not PVT-core) MSN synapses (H.M. Kuhn, S.J. Weber, E.-K. Hwang, M.M. Beutler, M.E. Wolf, Society for Neuroscience abstract 2024). The discrepancy between our results and the previous study (Alonso-Caraballo et al., 2024) may reflect differences in the oxycodone regimen, rat strain, area of PVT targeted with ChR2, duration of abstinence (>AD21 in our studies), or the fact that the previous study extracted brains for recordings 30-60 min after completion of the AD14 seeking test. For cocaine, a cue-induced seeking after incubation is associated with transient CP-AMPAR internalization (Wright et al., 2020). Studies in PVT-D1 MSN and PVT-A2a MSN after oxycodone incubation are underway.

### Opioid incubation and sex differences

Comparison of behavioral data for male and female rodents has yielded mixed results in different models of opioid addiction (Lopresti et al., 2020). However, our data are in line with other findings indicating that sex does not robustly influence incubation of opioid craving (Nicolas et al., 2022; Chow, 2025). Focusing first on the self-administration training period, we did not observe sex differences in active or inactive hole responding or oxycodone infusions, consistent with several other long-access studies [(de Guglielmo et al., 2019; Alonso-Caraballo et al., 2024; Patel and Loweth, 2024); also see (Hinds et al., 2023)], although modest sex differences in some parameters were found in others (Mavrikaki et al., 2017; Kimbrough et al., 2020; Mavrikaki et al., 2021; Guha et al., 2022).

We also observed no sex differences in incubation of oxycodone craving, assessed by comparing responding in the previously active hole on AD15 or AD30 to responding on AD1, consistent with many studies demonstrating oxycodone incubation in males (Blackwood et al., 2019b; Blackwood et al., 2019a; Bossert et al., 2019; Salisbury et al., 2020; Altshuler et al., 2021a; Altshuler et al., 2021b; Fredriksson et al., 2021b; Wong et al., 2022; Fredriksson et al., 2023; Patel and Loweth, 2024) and others demonstrating incubation in females (Fredriksson et al., 2020; Patel and Loweth, 2024). However, one study observed significant incubation in females but not males on AD14 (Alonso-Caraballo et al., 2024) and another found higher seeking on AD14 in females (Guha et al., 2022). One limitation of our study is that the effect of estrous cycle was not addressed. While recent studies found no effect of estrous cycle on oxycodone seeking after forced abstinence on AD15 (Olaniran 2023) or cue-induced reinstatement of oxycodone seeking after extinction (Hinds et al., 2023), another found that females in estrus at the time of an AD43-45 seeking test did not exhibit incubation whereas non-estrus females did (Patel and Loweth, 2024). Finally, no sex differences have been found for heroin incubation (Venniro et al., 2017; Venniro et al., 2019; Bossert et al., 2021; D’Ottavio et al., 2022; Mayberry et al., 2022b; Nett and LaLumiere, 2022) or morphine incubation (Mayberry et al., 2022a), and no effects of estrous cycle were seen for heroin incubation (D’Ottavio et al., 2022; Mayberry et al., 2022b).

Overall, then, the literature suggests that incubation of opioid craving is similar in male and female rats (Nicolas et al., 2022; Chow, 2025). It is difficult to assess whether this aligns with clinical data. While men and women differ in the path to opioid addiction and in patterns of use (Marsh et al., 2018; Becker and Chartoff, 2019), measures of craving and relapse during treatment or after long-term follow-up have indicated greater female vulnerability, greater male vulnerability, or no difference (Nicolas et al., 2022). However, we acknowledge that interpretation of such studies is complicated because most subjects received opioid agonist therapy, which will decrease craving in both sexes. This underscores the need to assess incubation of opioid craving in humans using an appropriate abstinence design (Chow, 2025).

It is important to note that other behaviors that accompany opioid incubation (Mayberry et al., 2022b) and transcriptomic data after incubation (Mayberry et al., 2022a) do show sex differences. The latter study suggests the possibilities of different mechanisms supporting opioid incubation in males and females. This does not seem to be the case for the CP-AMPAR plasticity that is our focus, although we cannot rule out the possibility of small sex differences that were not revealed with our sample sizes. Similarly, male and female rats did not appear to differ with respect to adaptations at PVT-NAc shell synapses associated with oxycodone incubation (Alonso-Caraballo et al., 2024).

## Conclusions

This study shows that, during incubation of oxycodone craving, CP-AMPARs accumulate in excitatory synapses on D1 and A2a MSN in both core and shell subregions. This explains the ability of CP-AMPAR blockade in either NAc core or shell to prevent expression of oxycodone incubation (Wong et al., 2022). Along with prior studies of cocaine and methamphetamine incubation, this work establishes a common role for CP-AMPAR upregulation in incubation of opioid and psychostimulant craving. Although more work is required to understand differences in mechanisms and pathway-specificity of CP-AMPAR plasticity between drug classes, this commonality may suggest that abstinence-triggered mechanisms (e.g., homeostatic plasticity or silent synapse filling) are important for driving CP-AMPAR plasticity more so than specific actions of a particular drug class during the period of self-administration. Overall, our model is that CP-AMPAR insertion strengthens NAc synapses, enabling them to respond more robustly to drug cues, which in turn promotes greater cue-induced drug seeking.

## Acknowledgements

We thank Alana Moutier and Jonathan Funke for help with drug self-administration experiments.

## Author contributions

KAM and MEW developed the project; KAM, HMK and MEW wrote the manuscript; KAM, HMK, EKH, and MMB conducted the experiments.

## Present address for Kimberley A. Mount

Division of Behavioral Health and Recovery, Washington State Health Care Authority

---

**Figure S1.**
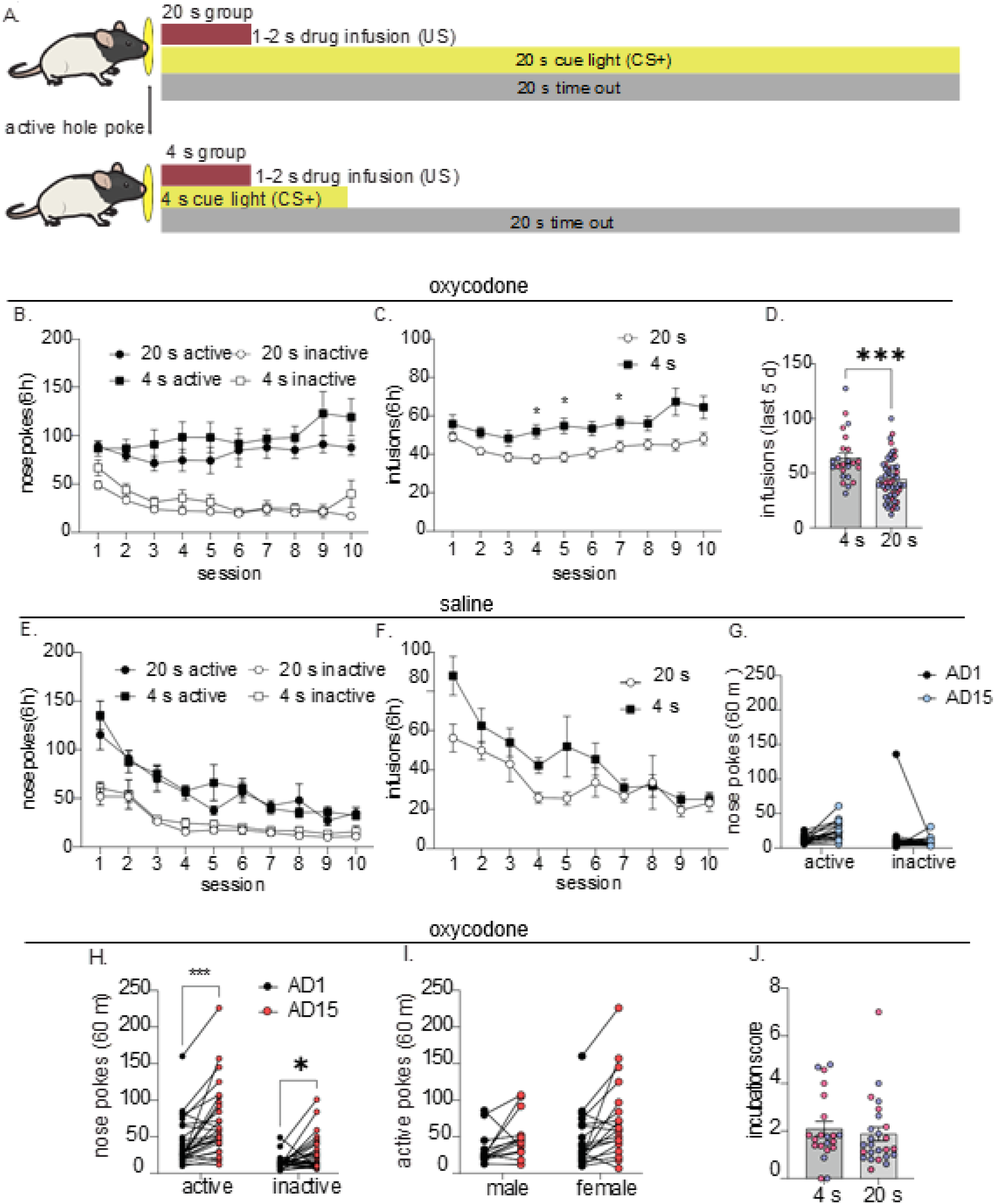
Shortening the cue light increases oxycodone intake but does not strengthen incubation. We experimented with shortening the cue light duration based on the possibility that the 20 second (s) light cue might to some degree become associated with the concurrent timeout period in which drug is not available rather than with the drug infusion itself, thereby attenuating cue-induced oxycodone seeking. (A) Diagram illustrating the 20 s cue light group (also shown in Figs. 1 and 2) and the 4 s cue light group. The 20 s group consisted of 15 saline rats (9 males, 6 females) and 58 oxycodone rats (36 males, 22 females). The 4 s group consisted of 19 saline rats (13 females, 6 males) and 33 oxycodone rats (19 females and 14 males). (B) The number of nose poke responses during oxycodone self-administration training was assessed using a mixed ANOVA with cue duration as the between subject factor, hole as within subject factor, and session as the repeated measure. We found a main effect of session (F(9,1588)=2.974, *p*=0.0016), hole (F(1,180)=81.12, *p*<0.0001), and a session x hole interaction (F(9,1588)=5.309, *p*<0.0001), but no main effect of cue duration (*p*>0.05). (C,D) Infusions during oxycodone self-administration were compared between the 4 s and 20 s groups using a mixed model ANOVA with cue duration as the between subjects factor and session as the repeated measure. Analysis revealed a main effect of session (F(9,791)=5.811, *p=*0.0007) and cue duration (F(1,90)=17.25, *p<*0.0001). Rats in the 4 s group took significantly more drug during sessions 4, 5 and 7 compared to rats in the 20-s group (all *p<*0.05; Fig. S1C) and significantly more drug across the last 5 days of training (unpaired *t*-test, *t(*80)=3.985, *p=*0.0001; Fig. S1D). There was no sex difference in oxycodone infusions in the 4 s group, as previously shown with the 20 s group (unpaired *t*-test, *p>*0.05, Fig. S1D; blue = males, magenta = females). In summary, the cue light duration did not affect the number of active or inactive nose pokes during oxycodone self-administration but the 4 s cue light was associated with greater oxycodone intake. Furthermore, the data indicate that rats trained with the 20 s cue made more active hole nose-pokes during the time out period than rats who received a 4 s cue, as evidenced by the same number of nose pokes but significantly fewer daily infusions in the 20 s group. (E,F) Using the same statistical approaches described above for the oxycodone rats, we found that nose pokes performed by saline rats also showed a main effect of session (F(9,658)=37.88, *p*<0.0001), hole (F(1,74)=47.80, *p*<0.0001) and a session x hole interaction (F(9,658)=4.012, *p*<0.0001) but no main effect of cue duration (*p*>0.05) (Fig. S1E). For saline infusions, we found a main effect of session (F(9,331)=11.76, p<0.0001), but not cue duration (Fig. S1F). (G) Saline rats (and oxycodone rats; see H) trained on a 4 s cue received 60 min cue-induced seeking tests on abstinence day (AD)1 and AD15. A mixed model ANOVA with treatment as the between subjects factor, hole as the within subjects factor, and AD as the repeated measure revealed no increase in active hole responses as a function of AD (*p*>0.05); furthermore saline rats did not discriminate between active and inactive holes during either seeking test (*p*>0.05). (H) Applying the same statistical analysis described for (G) to oxycodone rats trained on a 4 s cue revealed a main effect of AD (F(1,50)=14.82, *p*<0.0003), hole (F(1,44)=14.40, *p*=0.0004) and treatment (F(1,50)=24.98, *p*<0.0001) as well as interactions between AD x hole (F(1,44)=5.299, *p*=0.0261), AD x drug (F(1,44)=7.120, *p*=0.0106) and hole x treatment (F(1,44)=11.18, *p*=0.0017). Oxycodone rats showed significant increases in active hole responding on AD15 vs. AD1 (*p*=0.0142) and discriminated between the previously active and inactive holes on AD1 (*p*<0.0001) and AD15 (*p*<0.0001). (I) Comparison of the number of active hole responses on AD1 and AD15 between male and female rats in the 4 s oxycodone group using a mixed model ANOVA with sex as the between subject factor and AD as the repeated measure revealed a main effect of AD (F(1, 31)=14.04, *p*=0.0007) but not sex (*p*>0.05), although we did detect a moderate sex effect size (*d*=0.521). (J) To determine whether oxycodone rats in the 4 s group show stronger incubation compared to rats in the 20 s group, we compared the number of active hole nose pokes on AD1 and AD15 in each group. Because seeking test duration differed for these groups (60 min for the 4 s group and 30 min for the 20 s group), analysis was performed on the first 30 min of the test for the 4 s group. A two-way ANOVA with cue duration as the between subjects factor and AD as the repeated measure revealed a main effect of cue duration (F(1, 30)=12.66, *p*=0.0013), driven by significantly more active hole nose pokes on both AD1 (*p*=0.004) and AD15 (*p*=0.026) for rats in the 4 s group (data not shown). We further analyzed whether cue duration impacts the strength of incubation by comparing incubation scores (AD15 active pokes/AD1 active pokes) between the two cue duration groups. We did not detect a significant difference in incubation score (unpaired t-test, *p*>0.05) (Fig. 3J), although we observed that 86.6% of rats in the 4 s group had an incubation score >1 on AD15 compared to 76.9% in the 20 s group. In summary, shortening the light cue duration did not result in significantly stronger incubation of oxycodone craving. Since incubation did not differ between rats run under the two regimens, both types of rats were used for electrophysiological studies presented in Fig. 3.

**Figure S2.**
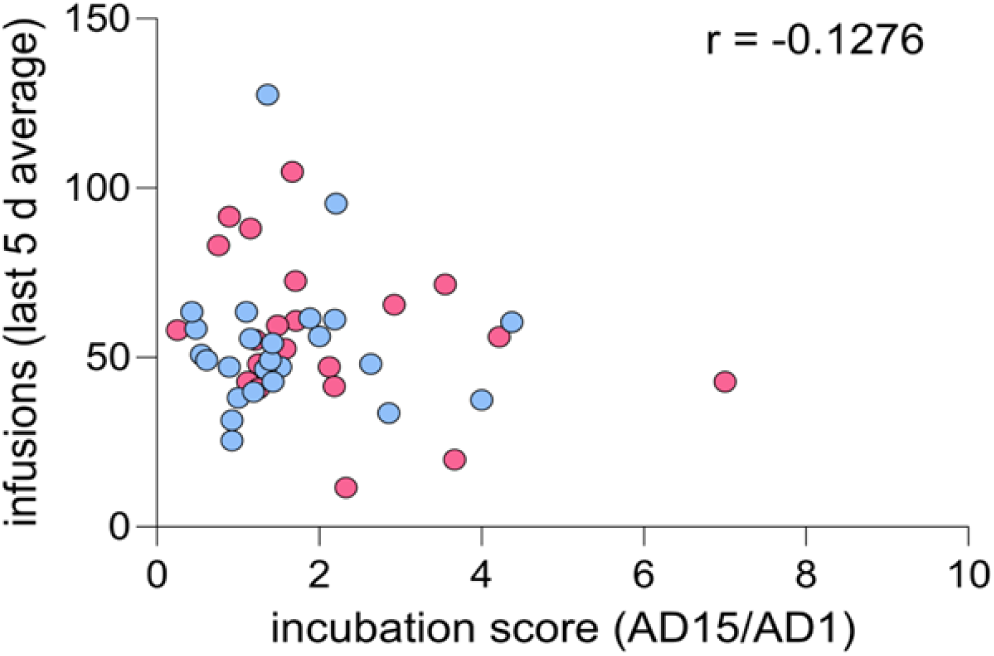
Rats express incubation and CP-AMPAR upregulation across a range of drug intake. An incubation score was calculated by dividing the number of active nose pokes on AD15 by the number of active nose pokes on AD1. There was no significant correlation between the incubation score and the average number of oxycodone infusions over the last 5 days of self-administration. Data are from WT Long-Evans rats shown in Figs. 2 and 3 (*n* = 21 rats; blue = males, magenta = females).

**Figure S3.**
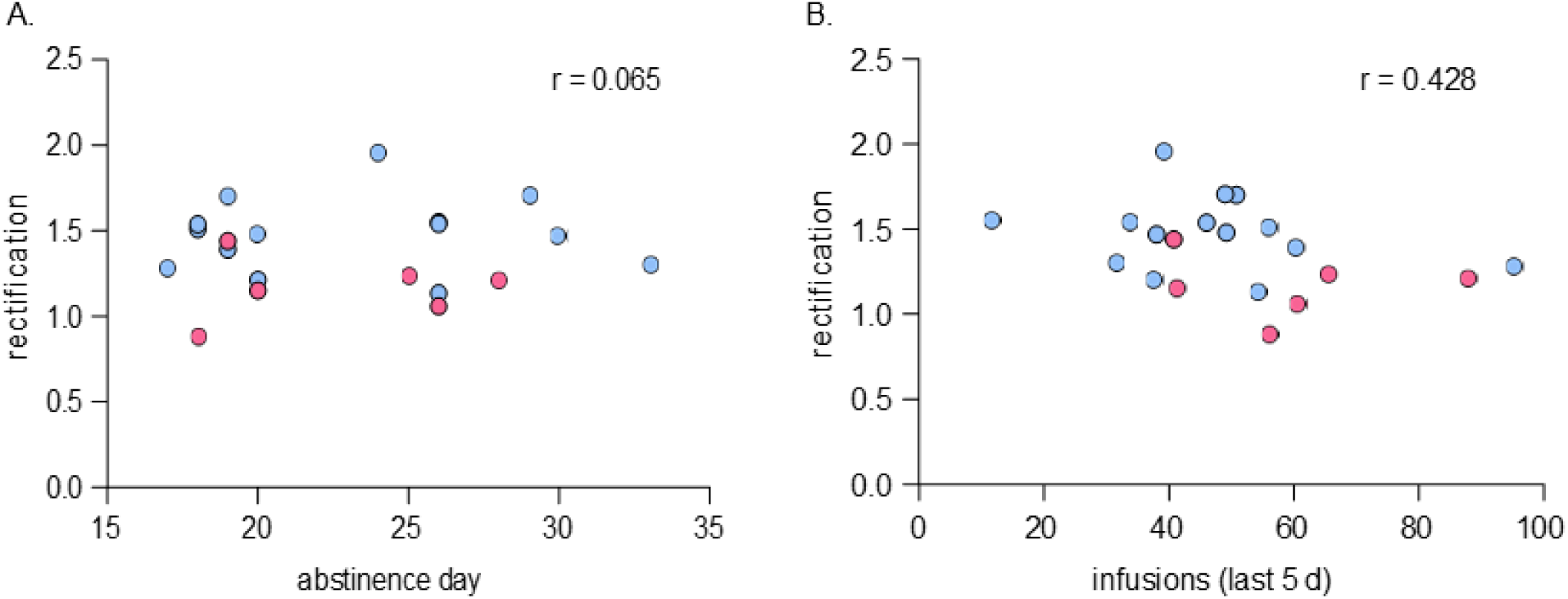
Rectification remains steady across abstinence days 17-33 and does not correlate with oxycodone intake during self-administration. (A) Correlation analysis evaluating the relationship between average rectification index (RI) from all cells per rat and length of days in abstinence from oxycodone self-administration at the time of recording (ranging between 17-33 days). No significant correlation was observed when NAc core and shell data were combined as shown in the figure (r=0.065, *p*>0.05) or when the subregions were analyzed separately (core r=-0.07, p=0.90; shell r=-0.46, p=0.29; data not shown). (B) Correlation analysis evaluating the relationship between the amount of oxycodone intake averaged across the last 5 days of self-administration and the average RI of all cells recorded from each rat. No significant correlation was observed when core and shell data were combined as shown in the figure (r=0.428, *p*>0.05) or when the subregions were analyzed separately (core: r=-0.68, p=0.14; shell: r=0.46, p=0.29; data not shown). Data are from WT Long-Evans rats shown in Figs. 2 and 3 (*n*=18 rats, 1-5 cells per rat; blue=males, magenta=females).

